## Supplemental Information for "Neural coding of choice and outcome are modulated by uncertainty in orbitofrontal but not secondary motor cortex"

#### Supplementary Figure 1

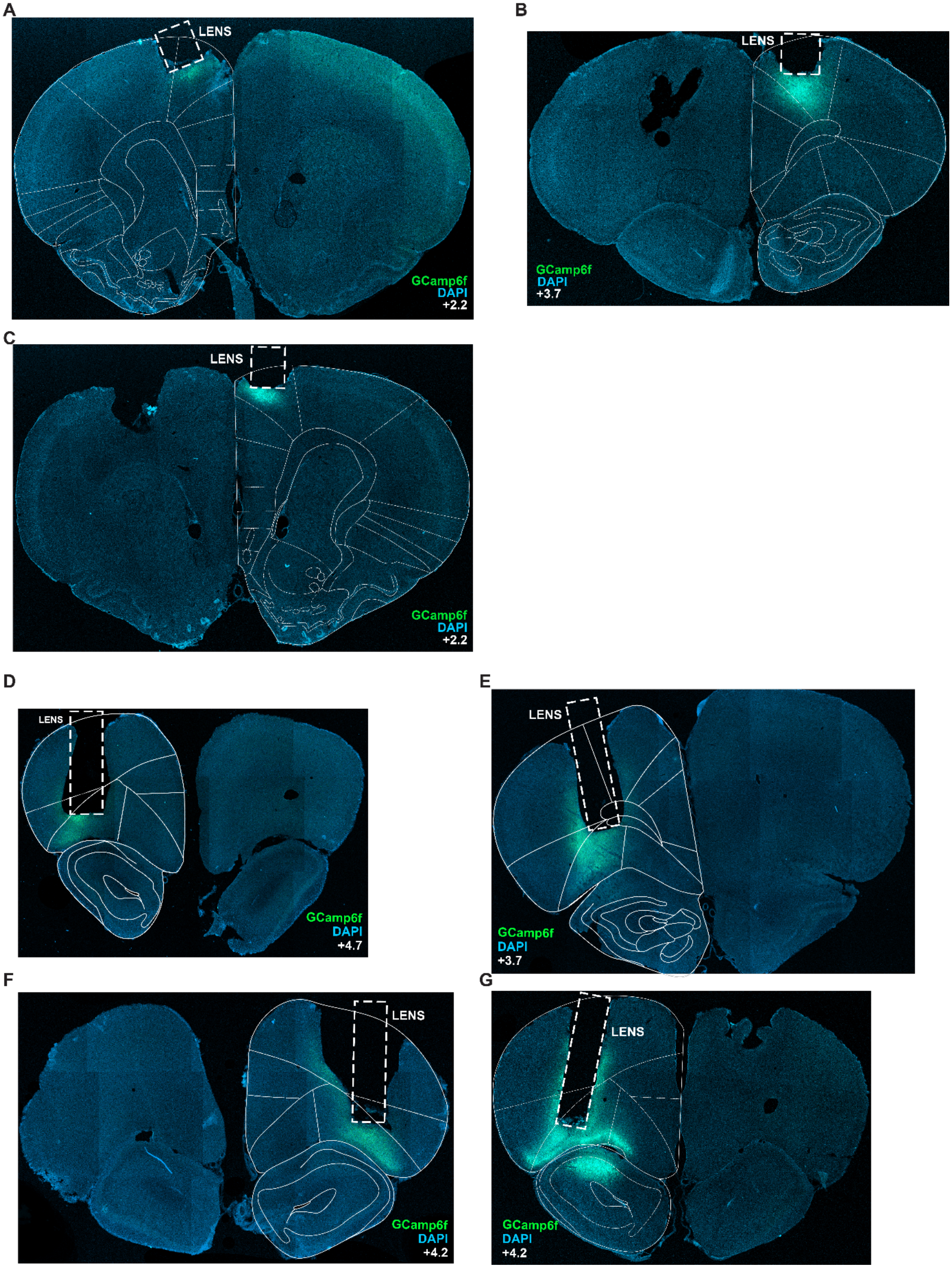


***Supplementary Figure 1.*** Photomicrographs of GCaMP6f and lens placement for all subjects included in the analyses. **(A-C)** Lenses in M2. **(D-G)** Lenses in OFC.

#### Supplementary Figure 2

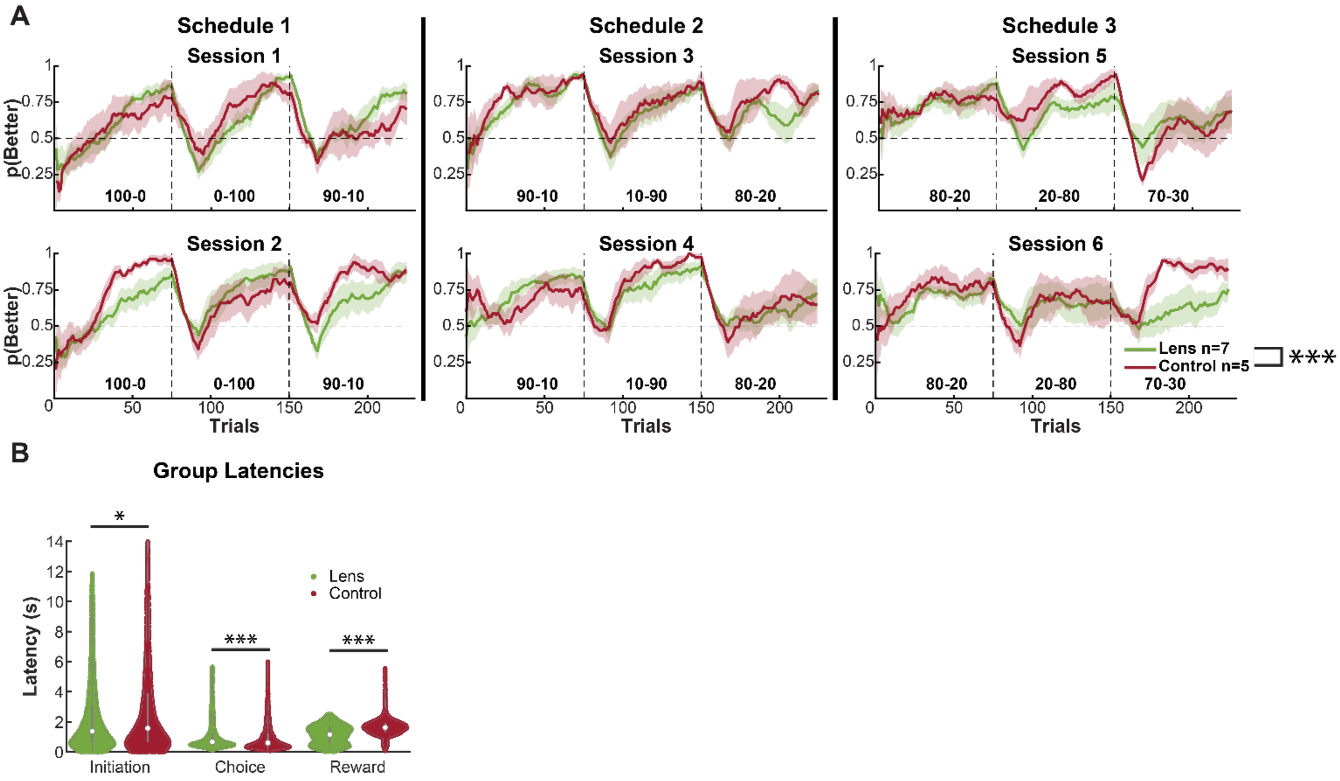


***Supplementary Figure 2.*** Learning in tethered-implanted rats is attenuated compared to untethered viral controls with no implants. **(A)** Tethered rats performed less accurately compared to viral non-tethered controls. **(B)** Controls initiated trials and collected rewards more quickly. However, tethered rats committed choices more quickly.

**Supplementary Figure 3**

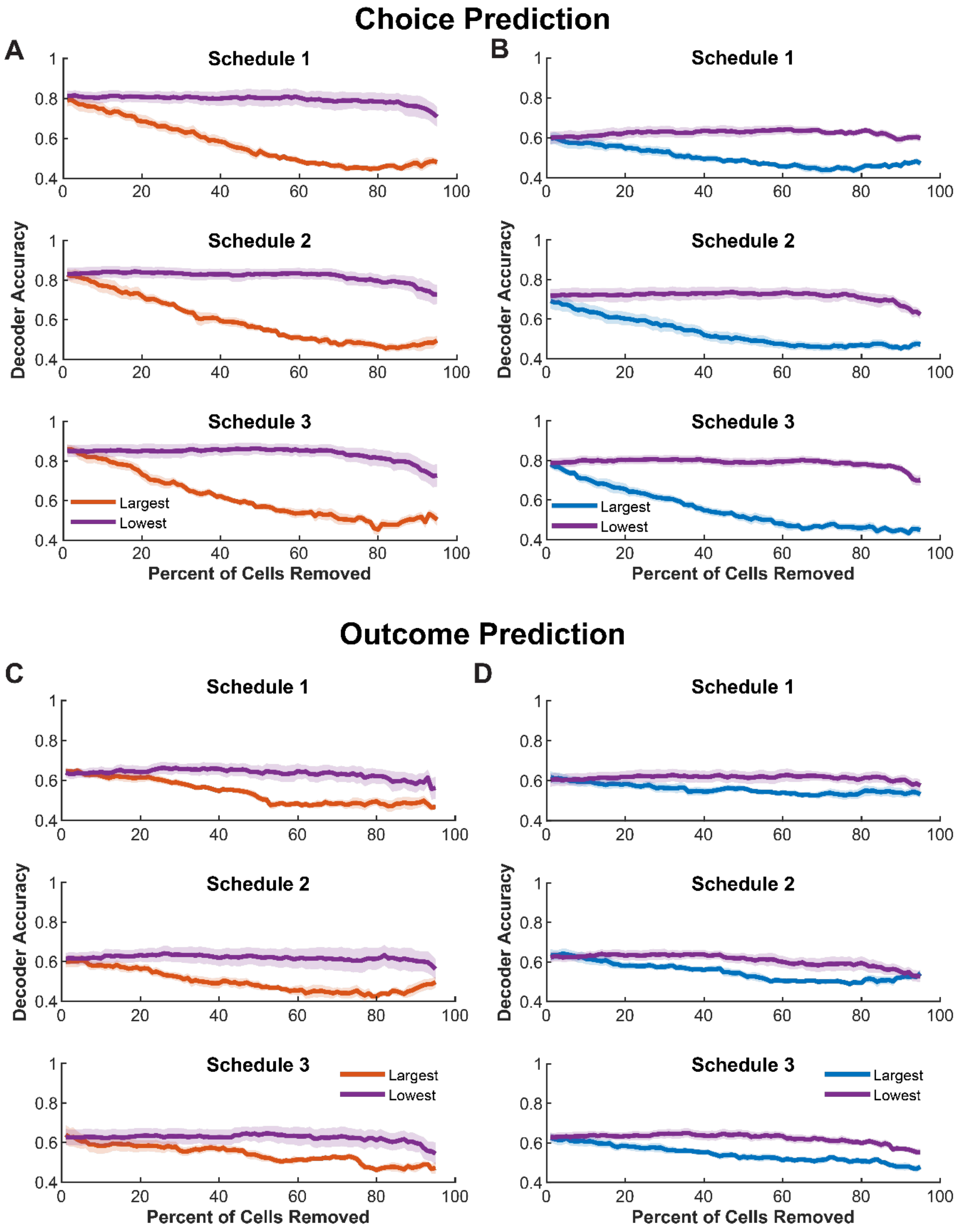


***Supplementary Figure 3. Removing neurons with the largest absolute beta coefficients affects decoder accuracy. (A)*** Decoder accuracy with an increasingly large percentage of neurons removed from the bin just prior to choice using neural data from M2. The purple line represented neurons with the lowest absolute beta coefficient from a GLM removed first and the orange line with the neurons that has the largest absolute beta coefficient removed first. **(B)** Same as A but with neural data from OFC. **(C)** Same as A but for Trial Outcome. **(D)** Same as B but for Trial Outcome.

**Strength and stability of encoding of trial types increases with uncertainty in OFC, but not M2**

We further probed differences in decoding accuracy of *Chosen Side* and *Trial Outcome* in M2 and OFC via a principal component analysis (PCA), which predicts neural correlates of behavior without behavioral measurements [1]. We applied this to neural data (*detrended F*) for the entire session and then selected a window of 2-s before and after the choice. When plotting this window using the first three principal components, we found that the trajectories diverged during the choice point and in other sessions, the separation occurred during the reward cue (**Supplementary Figure 4A-B**).

We first assessed the number of components necessary to explain 95% of the variance in a session and how these components changed from one schedule to the next. Since a different number of neurons was recorded per session, we limited the PCA to a randomly selected number of cells equal to the session with the lowest number of cells (49), resampling 500 times. We found a significant main effect on Area (GLM: $\beta_{Area}$ = -5.866, $p$ = 0.0077) and a significant Area x Schedule interaction (GLM: $\beta_{Area:Schedule}$ = 2.243, $p$ = 0.0254) (**Supplementary Figure 4C, Supplementary Table 5**), with only Schedule 1 (GLM: $\beta_{Area}$ = -2.968, $p$ = 0.0347) different by area (**Supplementary Table 6**): M2 greater than OFC in explained variance.

Due to the separation in 3-dimensional space we observed, we performed a Hotelling's T-squared distribution ($T^2$) analysis to test if there was a significant difference in the location in reduced space between the choice point on left/right trials and the reward cue during win/lose trials. Interestingly, for M2 we found that every session in Schedules 1 and 2 resulted in a significant separation between left and right trials at the choice point. In contrast, in OFC only about half of the sessions resulted in a significant separation during the first two schedules. During Schedule 3, the pattern was reversed with more significant separation for OFC than M2 (**Supplementary Figure 4D**). We observed a similar trend when analyzing the separation at the time of reward (**Supplementary Figure 4E**).

Finally, to further understand which aspect of the population dynamics drives the decoder results, we compared the coding of trial types in single neurons. We focused on the same time points used for our statistical analyses (before choice and reward cue) and selected cells using decoder weights that were 1 SD higher or lower from the mean at these time points and balanced the sampling such that there were equivalent left vs. right and rewarded vs. unrewarded (i.e., win vs. lose) trials. We then took the average activity of those cells for the specific trial types (*Chosen Side* vs. Trial Outcome) and computed correlations of the same trial type on odd and even trials (500 iterations of randomized sets of balanced trials), resulting in a set of correlation matrices (**Supplementary Figure 4F-G**). We compared these values to 500 times bootstrapped shuffled values. All values in M2 for the *Chosen Side* were significant throughout all schedules; however, only values in Schedules 2 and 3 were significant for OFC (**Supplementary Figure 4F**). In contrast, all values for *Trial Outcome* were significant in OFC across all sessions, but only the values in Schedules 1 and 2 were significant in M2 (**Supplementary Figure 4G**). An analysis on *Chosen Side* stability resulted in a main effect of Area (GLM: $\beta$Area = -0.227, p = 0.0438) (**Supplementary Table 7**), whereas an analysis of *Trial Outcome* stability resulted in Area (GLM: $\beta_{Area} = -0.239$, $p = 0.0266$) and Trial Type (GLM: $\beta_{TrialType} = 0.043$, $p < 0.0001$) differences, with the addition of a significant interaction of Area x Schedule (GLM: $\beta_{Area:Schedule} = 0.114$, $p = 0.0222$) (**Supplementary Table 8**). A post-hoc resulted in a significant effect of schedule only in OFC (GLM: $\beta_{Schedule} = 0.099$, $p = 0.0035$), but not M2 (**Supplementary Table 9**). Taken together, these results indicate an increase in the strength and stability of encoding of trial types with uncertainty in OFC, but not M2.

#### Supplementary Figure 4

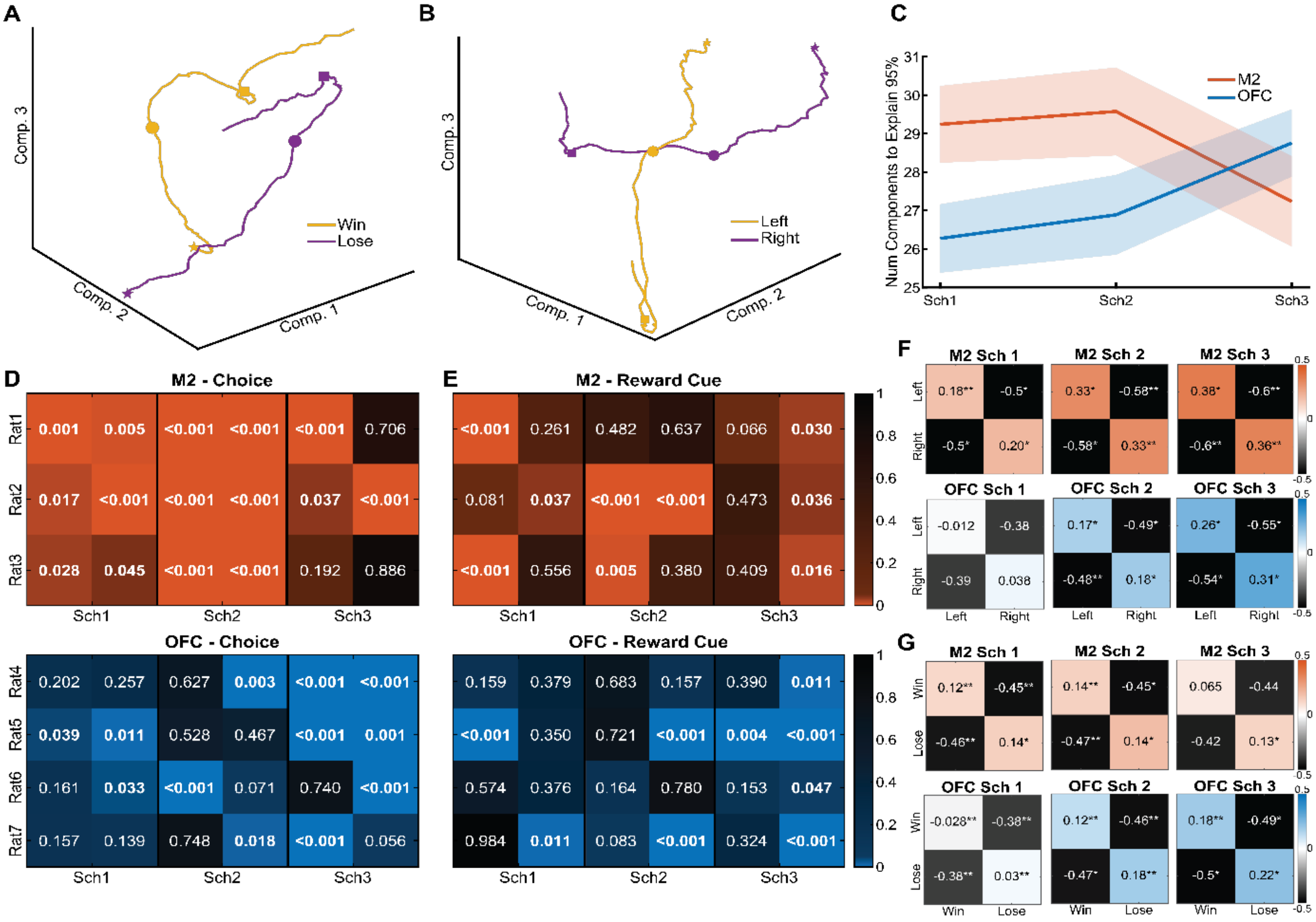


***Supplementary Figure 4. Differences in Neural Trajectories, Explained Variance, and Stability in M2 vs. OFC. (A)*** Example neural trajectory from M2. Shown are example trajectories with averaged coefficients for the first 3 principal components. STAR = beginning of a trial (2 s before choice), CIRCLE = choice, SQUARE = reward cue. In this example, the trajectories begin close to each other, move away during the choice, and return closer to each other during the reward cue. **(B)** Same as A, but with data from OFC. Instead of averaging across rewarded and unrewarded trials, we averaged across left and right trials. In this example, trajectories are close during the choice point but then move away for the reward cue. **(C)** Number of components needed to explain 95% of the variability across all schedules in M2 and OFC. We found a significant effect of Area and an Area x Schedule interaction, indicating a different pattern of explained variance in M2 and OFC. **(D)** Matrices of p-values from Hotelling's T2 to test if the choice point is significantly different between left and right trials across schedules. For M2, all values were below 0.05 during Schedules 1 and 2, only half were significant in Schedule 3. In contrast, Schedule 3 contained the most p-values below 0.05 in OFC. **(E)** Same as D but with the choice point on Reward Cue. There were significant sessions in each schedule in M2, whereas only Schedule 3 contained the greatest number of p-values below 0.05 in OFC. **(F)** Stability analysis for left vs right trials across sessions. An omnibus GLM resulted in a significant effect of Area only (Table 11). **(G)** Same as F but for rewarded (Win) and unrewarded (Lose) trials. An omnibus GLM resulted in a significant effect of Area and Trial Type. Additionally, there was a significant Area x Schedule interaction, with the post-hoc revealing an effect of schedule only in OFC, but not in M2. *p<0.05 **p<0.01 when comparing the results to shuffled data.

**M2 has a consistently larger ratio of Block Selective neurons compared to OFC**

M2 keeps track of different blocks better than OFC throughout the entirety of the trial. We assessed if single neurons in M2 and OFC were selective to blocks within a session **(Supplementary Figure 5A-C**), during the same epochs shown in **Figure 3**. For this analysis we included Chosen Side and Trial Outcome as covariates to remove all of the variance explained by these predictors and then tested if any of the remaining explained variance was significantly due to block. We found that M2 had a larger ratio of block selective neurons during choice prediction (GLM: $\beta_{Area} = -0.208$, $p = 0.0069$, **Supplementary Table 10**), outcome prediction (GLM: $\beta_{Area} = -0.2250$, $p = 0.0114$, **Supplementary Table 11**), and reward retrieval (GLM: $\beta_{Area} = -0.1757$, $p = 0.0242$, **Supplementary Table 12**).  As an example (**Supplementary Figure 5D**), this OFC cell is similarly active during Block 1 and 2 in choice prediction, but not during Block 3. Another example OFC cell is more active during Block 2 and Block 3, but not Block 1, in predicting outcome (**Supplementary Figure 5E**). And for the reward retrieval epoch, this example M2 cell is more active on Block 1, but then is not on Blocks 2 and 3 (**Supplementary Figure 5F**).

#### Supplementary Figure 5

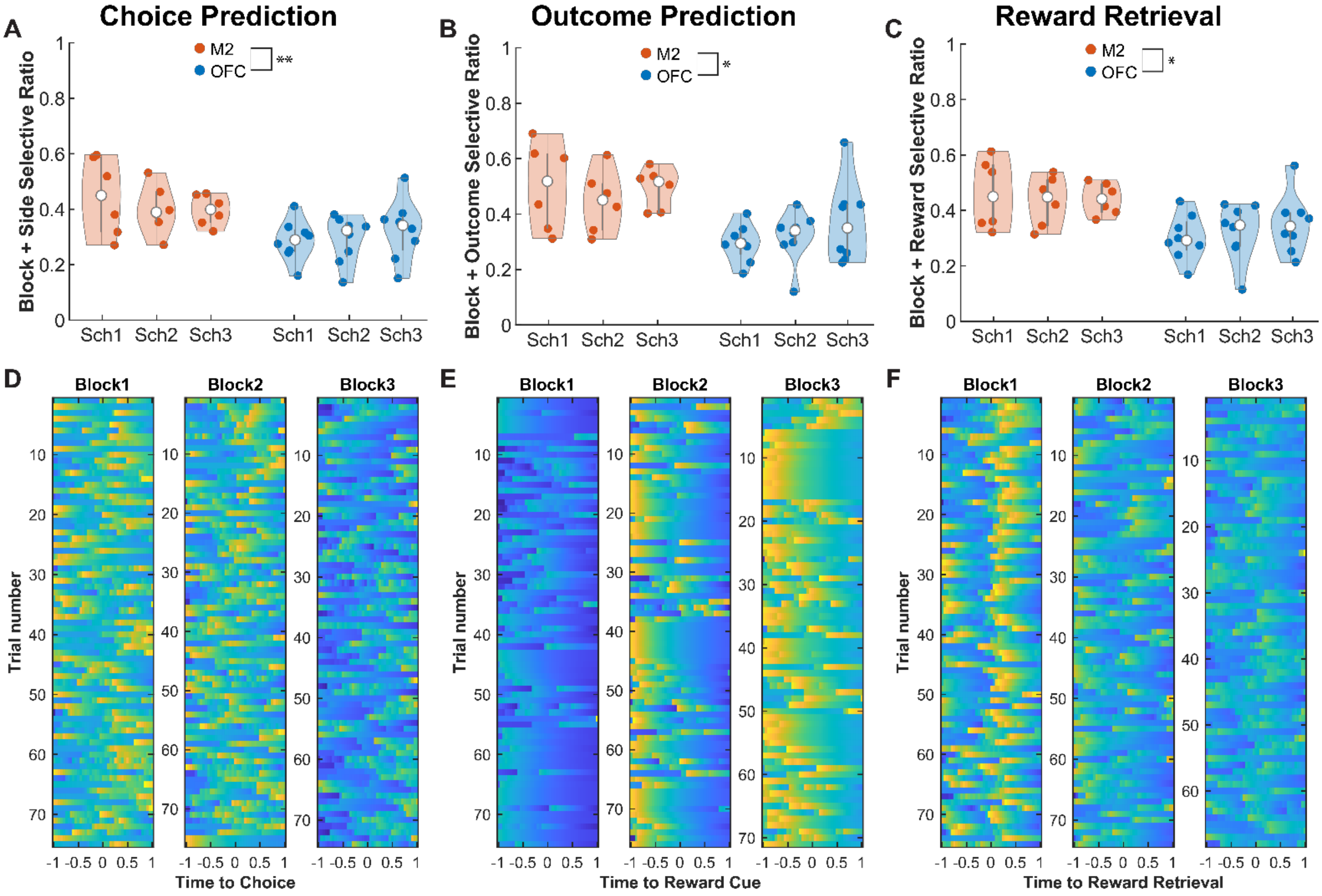


***Supplementary Figure 5. More Block Selective Neurons in M2 than OFC.*** **(A)** Ratio of block selective neurons during Choice Prediction is overall larger in M2 than OFC. **(B)** Same as A but during the Outcome Prediction epoch. **(C)** Same as A but during the Reward Retrieval epoch. **(D)** Example of the z-score normalized activity of a significant block selective OFC neuron on left and right trials during the Choice Prediction epoch. **(E)** Same as D but for a significant block-selective OFC cell during the Outcome Prediction epoch. **(F)** Same as D but for a significant block-selective M2 neuron during the Reward Retrieval epoch. *p<0.05. **p<0.01.

**Co-registration analysis reveals greater stability for Chosen Side Decoding**

Of recorded neurons, 29.4% of M2 and 44.7% of OFC neurons were co-registered between sessions 1 and 2 (**Supplementary Figure 6A**). We assessed if the co-registered neurons were selective for a specific trial variable (i.e., Chosen Side or Trial Outcome). Of all co-registered neurons in M2, 35.8% were not selective for any variable on any session, whereas 26.5% were selective for Chosen Side, 17.8% were selective for Trial Outcome, and 19.9% exhibited mixed selectivity. For OFC we found that 46.5% of these cells were non-selective, 22.8% were selective for Chosen Side, 17.6% were Trial Outcome-selective, and 10.9% exhibited mixed selectivity (**Supplementary Figure 6A**). We next investigated whether the ratio of cross-registered cells differed across schedule (**Supplementary Figure 6B**): We found a trend for a main effect of Schedule (GLM: $\beta_{Schedule}$ = -0.1793, $p$ = 0.0517, **Supplementary Table 13**), but no significant interaction of Area by Schedule (GLM: $\beta_{Area:Schedule}$ = 0.0847, $p$ = 0.1218, **Supplementary Table 13**). Finally, we wanted to assess if task variables such as Chosen Side and Trial Outcome if could be decoded only from co-registered cells from session 1 (training) to session 2 (testing). Decoding was better than chance for Chosen Side (**Supplementary Figure 6C**) but not for Trial Outcome (**Supplementary Figure 6D**). This indicates that M2 and OFC neurons exhibit greater stability for the encoding of the former compared to the latter.

### Supplementary Figure 6

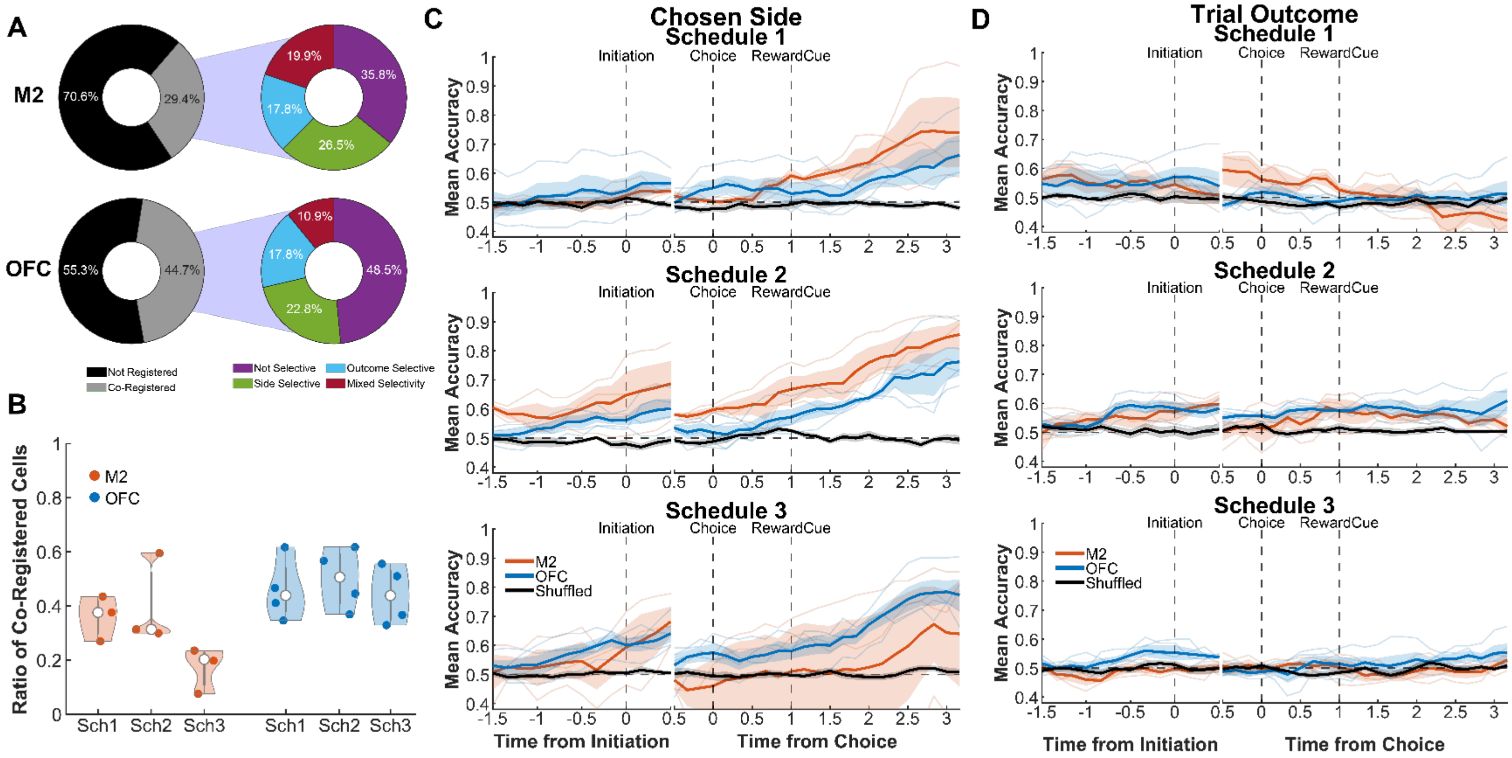


***Supplementary Figure 6. Co-registration analysis reveals greater stability for Chosen Side Decoding.*** **(A)** Left: Donut Chart indicating the percentage of all neurons imaged throughout all three of the schedules that were co-registered from the first-to-second session of that same schedule. Right: Donut chart indicating the selectivity of co-registered neurons. “Side selective” were neurons that exhibited significant phasic changes in the bin right before choice (same as Figure 3D). “Outcome selective” were neurons that exhibited significant phasic changes in the bin right before the reward cue (same as Figure 3E). “Mixed selectivity” were neurons that were selective for more than one type throughout the two sessions of each schedule. **(B)** Ratio of co-registered neurons per schedule for M2 and OFC. There were no significant differences. **(C)** Decoding accuracy for Chosen Side using only co-registered neurons. Training data came exclusively from session 1 and then was testing using trials from session 2. **(D)** Same as C but for Trial Outcome.

#### Supplementary Methods

*Principal Component Analysis*. Detrended fluorescence for the entire session was inputted into the pca function on Matlab with default parameters. Frames that had NaNs in them were interpolated if they were less than five continuous NaNs or if they were longer than five continuous frames, they were removed entirely from the dataset, its indices were saved, and after the PCA was finished, the same frames were filled with NaNs so that the results output would be the same length as the input. We saved the coefficients from the output, which contained the coefficients of all extracted components and the variability explained by each component. The example trajectories were plotted by selecting a 4-s window of the coefficients of the first three principal components centered around choice for every trial in a session. Then, win/lose or right/left trials were averaged, and the resulting 3D vector was plotted using the plot3 function in Matlab. Statistical comparisons and post-hocs for the explained variability were performed using Matlab's *fitglme* function and the formulas are included in the corresponding tables. For the *Chosen Side* comparisons in **Supplementary Figure 4D**, we selected all the coefficients of the first three principal components before choice and separated them according to left and right choice. Then, we used a custom function that performed a multivariate Hotelling's $T^2$, which tested the hypothesis that both groups came from the same population. For **Supplementary Figure 4E**, we did the same but separated every trial according to whether it was a win or a loss.

*Stability Analysis*. Neurons used in this analysis were selected for decoder weights that were either one standard deviation larger or smaller than the average weights used for cells in the decoder analysis. A 4-s window (1 s before and 3 s after) of detrended fluorescence around choice was obtained for every trial in a session and was temporally downsampled, similar to the decoder analysis. After downsampling trials were balanced, ensuring equal number of left/right and win/lose per session. The subset of trials that were used was randomly chosen, and the analysis was repeated 500 times. The trials were further split into odds and evens, and each cell's activity was averaged within their respective group. For example, in the *Chosen Side* analysis, there were four subgroups: left odd, left even, right odd, and right even, comprised of a cell-by-subgroup matrix for that analysis. Each group was then normalized using the *normalize* function with the 'range' method to compare across multiple cells, then *corrcoef* to correlate each group with another similar group. The resulting correlation matrix was stored and then averaged across the 500 samples. In parallel, the same analysis was performed but with shuffled data. Each result was compared with the shuffled equivalent using a t-test. Statistical comparisons and post-hocs were performed using Matlab's *fitglme* function, and the formulas are included in the corresponding tables.

*Block Selectvity Analysis*: Instead of fitting neural data from the epoch of choice for all trials in a session, here we separated the trials based on which block they came from. We also modified the formula used in the GLM: NeuralData ~ [1 + Block + ChosenSide + Rewarded], with Chosen Side and Rewarded as covariates. We designated neurons as 'Block selective,' if they were significant for Block.

*Cell Registration Analysis*: Cell Registration was performed using the Matlab CellReg package from the Ziv Lab [2], using the default setting and following the suggested parameters. Spatial footprints were loading into CellReg from the output in CaImAn. Cell registration was only performed from session 1 to session 2 within a specific schedule. Co-registered cell selectivity was categorized as "Chosen Side selective" if a cell exhibted significant phasic change in activity (decrease or increase) for Chosen Side at any point in the schedule. A cell was "Outcome selective" if it exhibited significant phasic change in activity for Reward/Unrewarded in the schedule. And finally, a cell was classified as exhibiting "Mixed selectivity" if it exhibited significant phasic change in activity for both both side and Rewarded/Unrewarded at any point in the schedule. The decoder analysis with co-registered cells was performed similarly to **Figure 2F-G** but only using co-registered cells, trained on session 1 and tested on session 2.

***Supplementary Table 1. Analysis of behavioral performance across all blocks in all sessions comparing lens-implanted with viral controls.***

| $\gamma$ = **Probability Better Option** | | | | | | | |
|---|---|---|---|---|---|---|---|
| Formula | $\gamma$~[1+Session*Block*Area+Cohort+(1+Session*Block*TrialNum\|RatID)] | | | | | | |
| Coefficients | **β** | SE | tStat | DF | P | CIL | CIU |
| Intercept | 1.1487 | 0.1436 | 8.0008 | 8091 | <0.0001 | 0.8673 | 1.4302 |
| **TrialNum** | 0.0104 | 0.0019 | 5.3930 | 8091 | **<0.0001** | 0.0066 | 0.0142 |
| Session | -0.0263 | 0.0387 | -0.6789 | 8091 | 0.4972 | -0.1021 | 0.0496 |
| **Area_Lens** | -0.1830 | 0.0537 | -3.4096 | 8091 | **0.0007** | -0.2882 | -0.0778 |
| **Block** | -0.7306 | 0.0938 | -7.7896 | 8091 | **<0.0001** | -0.9144 | -0.5467 |
| **TrialNum:Session** | -0.0012 | 0.0006 | -1.9805 | 8091 | **0.0477** | -0.0025 | 0.0000 |
| TrialNum:Block | -0.0003 | 0.0008 | -0.3164 | 8091 | 0.7517 | -0.0018 | 0.0013 |
| **Session:Block** | 0.0819 | 0.0345 | 2.3709 | 8091 | **0.0178** | 0.0142 | 0.1496 |
| TrialNum:Session:Block | 0.0000 | 0.0003 | 0.0634 | 8091 | 0.9495 | -0.0006 | 0.0006 |

Alpha Level 0.05.

***Supplementary Table 2. Analysis of initiation latency across all blocks in all sessions comparing lens-implanted with viral controls.***

| $\gamma$ = **Initiation Latency** | | | | | | | |
|---|---|---|---|---|---|---|---|
| Formula | $\gamma$~[1+Session*Block*TrialNum+Area+(1+Session*Block*TrialNum\|RatID)] | | | | | | |
| Coefficients | **β** | SE | tStat | DF | P | CIL | CIU |
| (Intercept) | 5.3143 | 1.8552 | 2.8645 | 9441 | 0.0042 | 1.6776 | 8.9510 |
| **TrialNum** | -0.0632 | 0.0128 | -4.9456 | 9441 | **<0.0001** | -0.0883 | -0.0382 |
| Session | -0.1312 | 0.2263 | -0.5798 | 9441 | 0.5621 | -0.5747 | 0.3123 |
| **Area_Lens** | -2.1790 | 0.9817 | -2.2196 | 9441 | **0.0265** | -4.1034 | -0.2546 |
| Block | 1.5195 | 0.8695 | 1.7475 | 9441 | 0.0806 | -0.1850 | 3.2239 |
| **TrialNum:Session** | 0.0109 | 0.0026 | 4.1473 | 9441 | **<0.0001** | 0.0058 | 0.0161 |
| **TrialNum:Block** | 0.0128 | 0.0049 | 2.6063 | 9441 | **0.0092** | 0.0032 | 0.0225 |
| Session:Block | -0.3313 | 0.2388 | -1.3874 | 9441 | 0.1654 | -0.7993 | 0.1368 |
| TrialNum:Session:Block | -0.0022 | 0.0018 | -1.2322 | 9441 | 0.2179 | -0.0056 | 0.0013 |

Alpha Level 0.05.

***Supplementary Table 3. Analysis of choice latency across all blocks in all sessions comparing lens-implanted with viral controls.***

| $\gamma$ = **Choice Latency** | | | | | | | |
|---|---|---|---|---|---|---|---|
| Formula | $\gamma$~[1+Session*Block*TrialNum+Area+(1+Session*Block*TrialNum\|RatID)] | | | | | | |
| Coefficients | **β** | SE | tStat | DF | P | CIL | CIU |
| (Intercept) | 2.0836 | 0.6253 | 3.3323 | 9441 | 0.0009 | 0.8579 | 3.3093 |
| TrialNum | -0.0169 | 0.0070 | -2.4129 | 9441 | 0.0158 | -0.0306 | -0.0032 |
| Session | -0.1392 | 0.0796 | -1.7486 | 9441 | 0.0804 | -0.2952 | 0.0168 |
| **Area_Lens** | 1.2993 | 0.3477 | 3.7373 | 9441 | **0.0002** | 0.6178 | 1.9808 |
| Block | 0.0520 | 0.3883 | 0.1339 | 9441 | 0.8935 | -0.7092 | 0.8132 |
| TrialNum:Session | 0.0016 | 0.0016 | 1.0247 | 9441 | 0.3055 | -0.0015 | 0.0047 |
| TrialNum:Block | 0.0043 | 0.0032 | 1.3602 | 9441 | 0.1738 | -0.0019 | 0.0106 |
| Session:Block | -0.0311 | 0.1157 | -0.2688 | 9441 | 0.7881 | -0.2578 | 0.1957 |
| TrialNum:Session:Block | -0.0004 | 0.0009 | -0.4092 | 9441 | 0.6824 | -0.0021 | 0.0014 |

Alpha Level 0.05.

***Supplementary Table 4. Analysis of reward latency across all blocks in all sessions comparing lens-implanted with viral controls.***

| $\gamma$ = **Reward Latency** | | | | | | | |
|---|---|---|---|---|---|---|---|
| Formula | $\gamma$~[1+Session*Block*TrialNum+Area+(1+Session*Block*TrialNum\|RatID)] | | | | | | |
| Coefficients | **β** | SE | tStat | DF | P | CIL | CIU |
| (Intercept) | 2.1173 | 0.2730 | 7.7547 | 6086 | <0.0001 | 1.5820 | 2.6525 |
| TrialNum | -0.0032 | 0.0039 | -0.8147 | 6086 | 0.4153 | -0.0109 | 0.0045 |
| Session | -0.1490 | 0.0845 | -1.7629 | 6086 | 0.0780 | -0.3147 | 0.0167 |
| **Area_Lens** | -0.7252 | 0.1534 | -4.7278 | 6086 | **<0.0001** | -1.0259 | -0.4245 |
| Block | -0.3421 | 0.2696 | -1.2690 | 6086 | 0.2045 | -0.8707 | 0.1864 |
| TrialNum:Session | 0.0011 | 0.0012 | 0.9228 | 6086 | 0.3561 | -0.0012 | 0.0034 |
| TrialNum:Block | 0.0021 | 0.0016 | 1.3160 | 6086 | 0.1882 | -0.0010 | 0.0053 |
| Session:Block | 0.1385 | 0.0904 | 1.5324 | 6086 | 0.1255 | -0.0387 | 0.3156 |
| TrialNum:Session:Block | -0.0009 | 0.0006 | -1.4887 | 6086 | 0.1366 | -0.0021 | 0.0003 |

Alpha Level 0.05.

***Supplementary Table 5. Analysis comparing components to explain 95% variability in all sessions.***

| $\gamma$ = **Number of Components** | | | | | | | |
|---|---|---|---|---|---|---|---|
| Formula | $\gamma$~[1+Area*Schedule+Session+(1\|RatID:Session)] | | | | | | |
| Coefficients | **β** | SE | tStat | DF | P | CIL | CIU |
| (Intercept) | 36.378 | 3.4497 | 10.545 | 37 | <0.0001 | 29.388 | 43.368 |
| Session | -0.3536 | 0.7783 | -0.4544 | 37 | 0.6522 | -1.9306 | 1.2234 |
| **Area** | -5.8660 | 2.0805 | -2.8195 | 37 | **0.0077** | -10.082 | -1.6505 |
| Schedule | -2.5387 | 2.2227 | -1.1421 | 37 | 0.2607 | -7.0424 | 1.9650 |
| **Area:Schedule** | 2.2434 | 0.9631 | 2.3294 | 37 | **0.0254** | 0.2920 | 4.1948 |

Alpha Level 0.05

***Supplementary Table 6. Post-Hoc Analysis comparing components to explain 95% variability in all sessions.***

| $\gamma$ = **Number of Components in Schedule 1** | | | | | | | |
|---|---|---|---|---|---|---|---|
| Formula | $\gamma$~[1+Area+(1\|RatID:Session)] | | | | | | |
| Coefficients | **β** | SE | tStat | DF | P | CIL | CIU |
| (Intercept) | 32.209 | 2.0532 | 15.687 | 12 | <0.0001 | 27.735 | 36.682 |
| **Area** | -2.9682 | 1.2463 | -2.3816 | 12 | **0.0347** | -5.684 | -0.2528 |
| $\gamma$ = **Number of Components in Schedule 2** | | | | | | | |
| Formula | $\gamma$~[1+Area+(1\|RatID:Session)] | | | | | | |
| Coefficients | **β** | SE | tStat | DF | P | CIL | CIU |
| (Intercept) | 32.263 | 2.3760 | 13.579 | 12 | <0.0001 | 27.086 | 37.440 |
| Area | -2.6879 | 1.4422 | -1.8638 | 12 | 0.0870 | -5.830 | 0.4543 |
| $\gamma$ = **Number of Components in Schedule 3** | | | | | | | |
| Formula | $\gamma$~[1+Area+(1\|RatID:Session)] | | | | | | |
| Coefficients | **β** | SE | tStat | DF | P | CIL | CIU |
| (Intercept) | 25.717 | 2.1901 | 11.743 | 12 | <0.0001 | 0.800 | 1.028 |
| Area | 1.5187 | 1.3293 | 1.1425 | 12 | 0.2755 | -0.131 | 0.007 |

Alpha Level 0.05.

***Supplementary Table 7. Analysis of chosen side coding stability***

| $\gamma$ = **Chosen Side Stability** | | | | | | | |
|---|---|---|---|---|---|---|---|
| Formula | $\gamma$~[1+Area*Schedule+Session+TrialType+(1\|RatID:Session)] | | | | | | |
| Coefficients | **β** | SE | tStat | DF | P | CIL | CIU |
| (Intercept) | 0.3003 | 0.1844 | 1.6287 | 78 | 0.1074 | -0.0668 | 0.6673 |
| Session | -0.0160 | 0.0414 | -0.3867 | 78 | 0.7000 | -0.0984 | 0.0664 |
| **Area** | -0.2268 | 0.1107 | -2.0497 | 78 | **0.0438** | -0.4471 | -0.0065 |
| Schedule | 0.0767 | 0.1182 | 0.6491 | 78 | 0.5182 | -0.1586 | 0.3121 |
| TrialType | 0.0219 | 0.0120 | 1.8176 | 78 | 0.0730 | -0.0021 | 0.0458 |
| Area:Schedule | 0.0446 | 0.0512 | 0.8703 | 78 | 0.3868 | -0.0574 | 0.1466 |

Alpha Level 0.05

***Supplementary Table 8. Analysis of trial outcome coding stability analysis.***

| $\gamma$ = **Trial Outcome Stability** | | | | | | | |
|---|---|---|---|---|---|---|---|
| Formula | $\gamma$~[1+Area*Schedule+Session+TrialType+(1\|RatID:Session)] | | | | | | |
| Coefficients | **β** | SE | tStat | DF | P | CIL | CIU |
| (Intercept) | 0.3097 | 0.1759 | 1.7612 | 78 | 0.0821 | -0.0404 | 0.6599 |
| Session | -0.0258 | 0.0395 | -0.6526 | 78 | 0.5159 | -0.1044 | 0.0529 |
| **Area** | -0.2387 | 0.1056 | -2.2601 | 78 | **0.0266** | -0.4489 | -0.0284 |
| Schedule | -0.0775 | 0.1128 | -0.6868 | 78 | 0.4942 | -0.3021 | 0.1471 |
| **TrialType** | 0.0482 | 0.0110 | 4.3875 | 78 | **<0.0001** | 0.0263 | 0.0701 |
| **Area:Schedule** | 0.1141 | 0.0489 | 2.3333 | 78 | **0.0222** | 0.0167 | 0.2114 |

Alpha Level 0.05

***Supplementary Table 9. Post-Hoc analysis comparing trial outcome stability across all sessions***

| $\gamma$ = **Trial Outcome Stability M2 Rats ONLY** | | | | | | | |
|---|---|---|---|---|---|---|---|
| Formula | $\gamma$~[1+Schedule+(1\|RatID:Session)] | | | | | | |
| Coefficients | **β** | SE | tStat | DF | P | CIL | CIU |
| (Intercept) | 0.1563 | 0.0802 | 1.9476 | 34 | 0.0598 | -0.0068 | 0.3193 |
| Schedule | -0.0150 | 0.0371 | -0.4035 | 34 | 0.6891 | -0.0905 | 0.0605 |
| $\gamma$ = **Trial Outcome Stability OFC Rats ONLY** | | | | | | | |
| Formula | $\gamma$~[1+Schedule+(1\|RatID:Session)] | | | | | | |
| Coefficients | **β** | SE | tStat | DF | P | CIL | CIU |
| (Intercept) | -0.0824 | 0.0695 | -1.1858 | 46 | 0.2418 | -0.2222 | 0.0575 |
| **Schedule** | 0.0991 | 0.0322 | 3.0804 | 46 | **0.0035** | 0.0343 | 0.1638 |

Alpha Level 0.05

***Supplementary Table 10. Analysis comparing block selective ratios during choice prediction epoch***

| $\gamma$ = **Ratio of Block Selective Neurons** | | | | | | | |
|---|---|---|---|---|---|---|---|
| Formula | $\gamma$~[1+Area*Schedule+Session +(1\|RatID)] | | | | | | |
| Coefficients | **β** | SE | tStat | DF | P | CIL | CIU |
| (Intercept) | 0.6635 | 0.1204 | 5.5086 | 37 | <0.0001 | 0.4194 | 0.9075 |
| Session | -0.0215 | 0.0256 | -0.8423 | 37 | 0.4050 | -0.0733 | 0.0302 |
| Schedule | -0.0283 | 0.0730 | -0.3872 | 37 | 0.7008 | -0.1761 | 0.1196 |
| **Area** | -0.2077 | 0.0727 | -2.8569 | 37 | **0.0070** | -0.3550 | -0.0604 |
| Area:Schedule | 0.0458 | 0.0316 | 1.4487 | 37 | 0.1559 | -0.0183 | 0.1099 |

Alpha Level 0.05

***Supplementary Table 11. Analysis comparing block selective ratios during outcome prediction epoch.***

| $\gamma$ = **Ratio of Block Selective Neurons** | | | | | | | |
|---|---|---|---|---|---|---|---|
| Formula | $\gamma$~[1+Area*Schedule+Session +(1\|RatID)] | | | | | | |
| Coefficients | **β** | SE | tStat | DF | P | CIL | CIU |
| (Intercept) | 0.7070 | 0.1400 | 5.0502 | 37 | <0.0001 | 0.4234 | 0.9907 |
| Session | 0.0002 | 0.0296 | 0.0057 | 37 | 0.9955 | -0.0599 | 0.0602 |
| Schedule | -0.0404 | 0.0846 | -0.4773 | 37 | 0.6359 | -0.2118 | 0.1310 |
| **Area** | -0.2250 | 0.0845 | -2.6622 | 37 | **0.0114** | -0.3962 | -0.0537 |
| Area:Schedule | 0.0380 | 0.0367 | 1.0363 | 37 | 0.3068 | -0.0363 | 0.1123 |

Alpha Level 0.05

***Supplementary Table 12. Analysis comparing block selective ratios during reward retrieval epoch.***

| $\gamma$ = **Ratio of Block Selective Neurons** | | | | | | | |
|---|---|---|---|---|---|---|---|
| Formula | $\gamma$~[1+Area*Schedule+Session +(1\|RatID)] | | | | | | |
| Coefficients | **β** | SE | tStat | DF | P | CIL | CIU |
| (Intercept) | 0.6471 | 0.1240 | 5.2199 | 37 | <0.0001 | 0.3959 | 0.8983 |
| Session | 0.0216 | 0.0271 | 0.7974 | 37 | 0.4303 | -0.0333 | 0.0765 |
| Schedule | -0.0811 | 0.0774 | -1.0482 | 37 | 0.3014 | -0.2379 | 0.0757 |
| **Area** | -0.1757 | 0.0748 | -2.3492 | 37 | **0.0243** | -0.3273 | -0.0242 |
| Area:Schedule | 0.0305 | 0.0335 | 0.9090 | 37 | 0.3692 | -0.0375 | 0.0984 |

Alpha Level 0.05

***Supplementary Table 13. Analysis comparing the ratio of co-registered neurons in all schedules.***

| $\gamma$ = **Ratio of Co-Registered neurons** | | | | | | | |
|---|---|---|---|---|---|---|---|
| Formula | $\gamma$~[1+Area*Schedule +(1\|RatID)] | | | | | | |
| Coefficients | **β** | SE | tStat | DF | P | CIL | CIU |
| (Intercept) | 0.5144 | 0.1945 | 2.6452 | 17 | 0.0170 | 0.1041 | 0.9247 |
| Schedule | -0.1793 | 0.0857 | -2.0928 | 17 | 0.0517 | -0.3600 | 0.0015 |
| Area | -0.0139 | 0.1180 | -0.1177 | 17 | 0.9077 | -0.2629 | 0.2351 |
| Area:Schedule | 0.0847 | 0.0520 | 1.6285 | 17 | 0.1218 | -0.0250 | 0.1944 |

Alpha Level 0.05

#### REFERENCES


1. Rubin, A., et al., *Revealing neural correlates of behavior without behavioral measurements*. Nature Communications, 2019. **10**(1): p. 4745.
2. Sheintuch, L., et al., *Tracking the Same Neurons across Multiple Days in Ca(2+) Imaging Data*. Cell Rep, 2017. **21**(4): p. 1102-1115.
